# Degradation of Red Blood Cell Deformability during Cold Storage in Blood Bags

**DOI:** 10.1101/2021.07.20.452409

**Authors:** Emel Islamzada, Kerryn Matthews, Erik Lamoureux, Simon P. Duffy, Mark D. Scott, Hongshen Ma

**Affiliations:** Department of Pathology and Laboratory Medicine, University of British Columbia; Centre for Blood Research, University of British Columbia; Department of Mechanical Engineering, University of British Columbia; British Columbia Institute of Technology; Canadian Blood Services; School of Biomedical Engineering, University of British Columbia; Vancouver Prostate Centre, Vancouver General Hospital

**Keywords:** Red blood cell, storage lesion, deformability, blood banking

## Abstract

Red blood cells (RBCs) stored in blood bags develop a storage lesion that include structural, metabolic, and morphologic transformations resulting in a progressive loss of RBC deformability. The speed of RBC deformability loss is donor-dependent, which if properly characterized, could be used as a biomarker to select high-quality RBC units for sensitive recipients or to provide customized storage timelines depending on the donor. We used the microfluidic ratchet device to measure the deformability of red blood cells stored in blood bags every 14 days over a span of 56 days. We observed that storage in blood bags generally prevented RBC deformability loss over the current standard 42-day storage window. However, between 42 and 56 days, the deformability loss profile varied dramatically between donors. In particular, we observed accelerated RBC deformability loss for a majority of male donors, but for none of the female donors. Together, our results suggest that RBC deformability loss could be used to screen for donors who can provide stable RBCs for sensitive transfusion recipients or to identify donors capable of providing RBCs that could be stored for longer than the current 42-day expiration window.

## INTRODUCTION

Red blood cells (RBCs) collected from donors for use in blood transfusions are currently stored at 4 °C for up to 42 days^1,2^. During this period, RBCs can develop a storage lesion, which is characterized by a number of structural (lipid peroxidation, Band 3 aggregation, membrane asymmetry), metabolic (slowed metabolism due to ATP and 2,3-diphosphoglycerate depletion), and morphologic transformations (discoid, echinocyte, and spherocyte)^3–6^. The storage lesion coincides with a shorter RBC circulation time arising from the rapid uptake of transfused RBCs by reticuloendothelial macrophages^7^, and thus resulting in the need for more frequent transfusions. While the 42-day storage window is currently uniformly applied to all RBC units, the actual rate of RBC degradation is known to vary between donors^4,5,8,9^. This variability has also been observed in outcomes for chronic transfusion recipients, where some RBC units are able to maintain durable hemoglobin levels in recipients; while other units are rapidly cleared, leading to the need for repeat transfusions^10,11^. Therefore, if the rates of degradation could be established for individual donors, it may be possible to select long-lasting units for sensitive recipients, such as those requiring chronic transfusions. Similarly, it may also be possible to provide customized expiration timelines for different donors to ensure that high-quality RBC units are not prematurely outdated, while less stable RBC units are used before they cease to provide clinical benefits. A key challenge in transfusion medicine has therefore been the development of a simple biomarker to assess the quality of stored blood to optimally meet the needs of the transfusion recipient.

Independent investigation of the cellular changes associated with RBC storage lesions has so far failed to produce a reliable biomarker for storage based degradation^12^. However, these cellular changes collectively reduce RBC deformability and thus makes this parameter an attractive potential biomarker for the RBC storage lesion. Previous studies have found the deformability of cold stored RBCs to be relatively stable for the first 14 days, but begins to degrade after 3 weeks of storage^5,13–16^. This change coincides with clinical evidence indicating that blood transfusion efficacy diminishes markedly after 30 days storage^17,18^. The loss of RBC deformability may directly impact on transfusion efficacy as more rigid RBC may be taken up more rapidly by the reticuloendothelial macrophages^19^. Additionally, rigid transfused RBCs are known to compromise microvascular flow by occluding blood capillaries^20^. Together, these findings from previous studies suggest that RBC deformability is a promising biomarker for the degradation of stored RBC units.

Various methods have been employed to measure deformability of stored RBCs, including bulk flow and single cell techniques. Bulk flow methods include micropore filtration^21,13,22^ and ektacytometry^23–25^. These methods infer RBC deformability indirectly based on blood viscosity and only provide a populational average measurement, both of which limits the sensitivity of these methods. Single cells methods, such as micropipette aspiration^26–28^ and optical tweezers^29–31^, provide single-cell deformability measurements, but are typically limited by sample throughput, which make them susceptible to variability and selection bias. Microfluidic techniques have been developed to overcome these limitations by enabling RBC deformability measurement with greater throughput and ease-of-use. Importantly, recent advances are beginning to provide sufficient sensitivity and repeatability to observed the loss of deformability in donated RBCs for use in blood transfusions^32–39^. These studies are beginning to suggest that it may be possible to identify blood donors that can provide high-quality RBC units that could be reserved for sensitive or chronic transfusion recipients. Donor-specific RBC deformability measurement is particularly useful for its potential to explain evidence for donor-dependent storage and transfusion efficacy^1,40–43^. Therefore, measurement sensitivity and repeatability are critical properties in efforts to assess differences between donated RBC units or between donors.

Recently, we developed a microfluidic technology to measure RBC deformability with sufficient sensitivity and repeatability for analyzing differences between healthy donors^36^. Using an accelerated aging model of RBCs stored in plastic tubes, we found that donor RBCs had degradation profiles that were highly variable between donors, but consistent for each donor. Importantly, some donors showed significant loss of RBC deformability during storage, while other donors showed little or no storage induced loss of RBC deformability. Here, to evaluate variability between donors during cold storage in blood bags, we assessed the degradation of RBC deformability over the 42-day storage window, and for an additional 14 days thereafter, for a total of 56 days. We show that, in most cases, blood bags preserved RBC deformability during the 42-day storage window, while the degradation of RBC deformability in the subsequent 14 days were highly variable. Our results confirm that RBC deformability provides a potentially useful approach for donor-level screening to identify donors for whom the storage expiration window could potentially be lengthened.

## METHODS

### Blood bags

This study was approved by the University of British Columbia’s Clinical Research Ethics Board (UBC REB# H19-01121) and Canadian Blood Services Research Ethics Board (CBS REB# 2019-029). Packed RBCs in standard Fresenius blood bags were collected and processed by Canadian Blood Services between January 2020 and February 2021.

### RBC storage and processing for deformability assessment

Packed RBC units were stored according to Canadian Blood Services (CBS) standard operating procedures, at 4°C for a period of 8 weeks, 2 weeks longer than the CBS-approved storage period of 42 days (6 weeks). Samples were analyzed on the day of RBC unit collection and processing, followed by analysis at weeks 2, 4, 6, and post-expiration at week 8.

To analyze the packed RBCs within a blood bag, a 3 ml sample was aseptically drawn from the unit through the blood administration ports, using a 27-gauge needle and syringe (BD). The ports were then covered with Parafilm^®^ to preserve the sterility of the unit throughout the storage period. The drawn sample was centrifuged at 1500 x g with no brakes for 10 min at room temperature, and supernatant was transferred to a fresh tube. The supernatant was centrifuged again at 1500 x g for 10 min to remove any remaining RBCs, transferred to a cryogenic vial and stored at −80°C for assessment of hemolysis at a later stage. Additionally, 100 μl of the blood drawn on day 1 was also stored for hemolysis assessment. The RBC pellet was suspended in Hank’s Balanced Salt Solution (HBSS, Gibco) and 0.2% Pluronic (F127, MilliporeSigma), and washed three more times, each time centrifuging for 5 minutes at 300 x g with brakes on. The final RBC sample was suspended at 1% hematocrit in HBSS+0.2% Pluronic solution, and used for deformability assessment using the Microfluidic Ratchet Device.

### Hematological parameters

At each sampling timepoint, 100 μl of packed RBC sample (no washing) was used to monitor the hematological parameter changes. Mean corpuscular volume (MCV), red blood cell distribution width (RDW-CV), mean cell hemoglobin (MCH), and mean corpuscular hemoglobin concentration (MCHC) was assessed using the Sysmex^®^ system.

### Assessment of hemolysis

The hemolysis assessment was performed as previously described^44^. Briefly, frozen whole blood and supernatant samples were fully thawed. An aliquot of each supernatant sample was transferred to a fresh tube and centrifuged at 3000 x g for 3 minutes. Packed, unwashed blood samples from day 1 of storage were vortexed for 30 seconds, and diluted 1:10 with DI water. Finally, 10 μl of each sample was transferred to a 96-well flat bottom plate (BioVision Inc) together with 100ul of Drabkin’s reagent (MilliporeSigma) containing Brij-35 solution (Thermofisher). The plate was then incubated at room temperature for 15 min on a plate shaker, and absorbance was read on a microplate reader (manufacturer) at 450 nm. Hemolysis was calculated using the following formula:

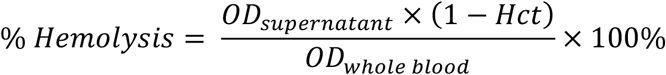

where *Hct* is the hematocrit, and *OD_supernatant_* and *OD_whole blood_* are the measured optical density from the supernatant and whole blood respectively.

### Microfluidic device manufacturing and operation

The Microfluidic Ratchet Device was manufactured as previously described^36,45,46^. Briefly, a mask with device features was created by photolithographic fabrication, which in turn was used to create a Polydimethylsiloxane(PDMS, Sylgard-184, Ellsworth Adhesives) master. The PDMS master device was used to create secondary molds for routine device manufacture. The PDMS device is made by mixing PDMS in a 10:1 ratio of PDMS and hardener and cured in the mold in a 65 °C oven for a minimum of 2 hours. Holes are manually punched in the device using Harris Uni-Core punches with 0.5 mm diameter for inlets and 2 mm for outlets. The PDMS part of the device is then bound to a thin RTV layer (RTV 615, Momentive Performance Materials LLC), followed by to a glass slide (2×3 inch, Corning) for durability, using a Harrick Plasma model PDC-001 air plasma.

Prior to sample introduction, the device is filled with PBSS+0.2% Pluronic buffer for 15 minutes until fully buffered. The microfluidic ratchet device operates using an oscillatory sorting pressure and constant forward pressure system, which propagates the sample forward towards the distinct outlet based on the cell’s ability to deform through the sorting matrix. The upward pressure is applied at 175 mbar for 4 seconds, and downward pressure is applied for 1 second at 162 mbar. The forward and sample pressures are applied at 40-45 and 50-55 mbar respectively. The distribution of cells in each distinct outlet is counted in each outlet microchannel. Each sample is run on 2 separate devices and the mean is calculated thereafter. Each device is discarded after each use.

### Statistical analysis

Statistical analysis was performed using GraphPad Prism (V8.0) software. Means and standard deviation from mean are plotted unless otherwise stated. To calculate the standard deviation for rigidity score obtained from doublet measurements, the following formula was used:

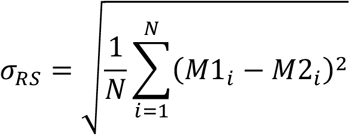

where M1 and M2 are the first and second RS measurements. Correlations between data sets were calculated using Pearson r with 95% confidence interval.

## RESULTS

### Sorting RBCs based on deformability using the microfluidic ratchet device

The design of the microfluidic ratchet device to sort RBCs based on deformability has been described previously^36,45,46^. Briefly, RBCs are deformed through a series of micrometer-sized constrictions using oscillatory flow, which selectively transport cells based on their ability to squeeze through each constriction. The constrictions are arranged in a matrix, where the openings of the constrictions are varied from 7.5 μm down to 1.5 μm, between rows in the matrix. This configuration enables sorting of RBCs based on their deformability into 12 fractions in different outlets. RBCs are sorted diagonally through the constriction matrix until reaching a limiting constriction row that prevents their transit. The RBCs then proceed horizontally along the limiting row of constrictions until they reach a specific outlet. The distribution of cells after sorting could be determined by imaging the flow of cells into the outlets or by counting the cells in the outlet via microscopy.

### Data Analysis

After sorting each RBC sample using the microfluidic ratchet device, the distribution of RBCs in outlets 1–12 can be used to establish a cumulative distribution from the smallest outlet to the largest outlet. The cumulative distribution could then be described using the rigidity score (RS) based on the outlet where the cumulative distribution function crosses 50% (**Fig. 1A-B**). Fractional outlet numbers can be obtained by linear interpolation of cumulative distribution graph between outlets. The RS provides a simple metric for comparing distributions between different donor and samples^36^.

**Fig. 1.**
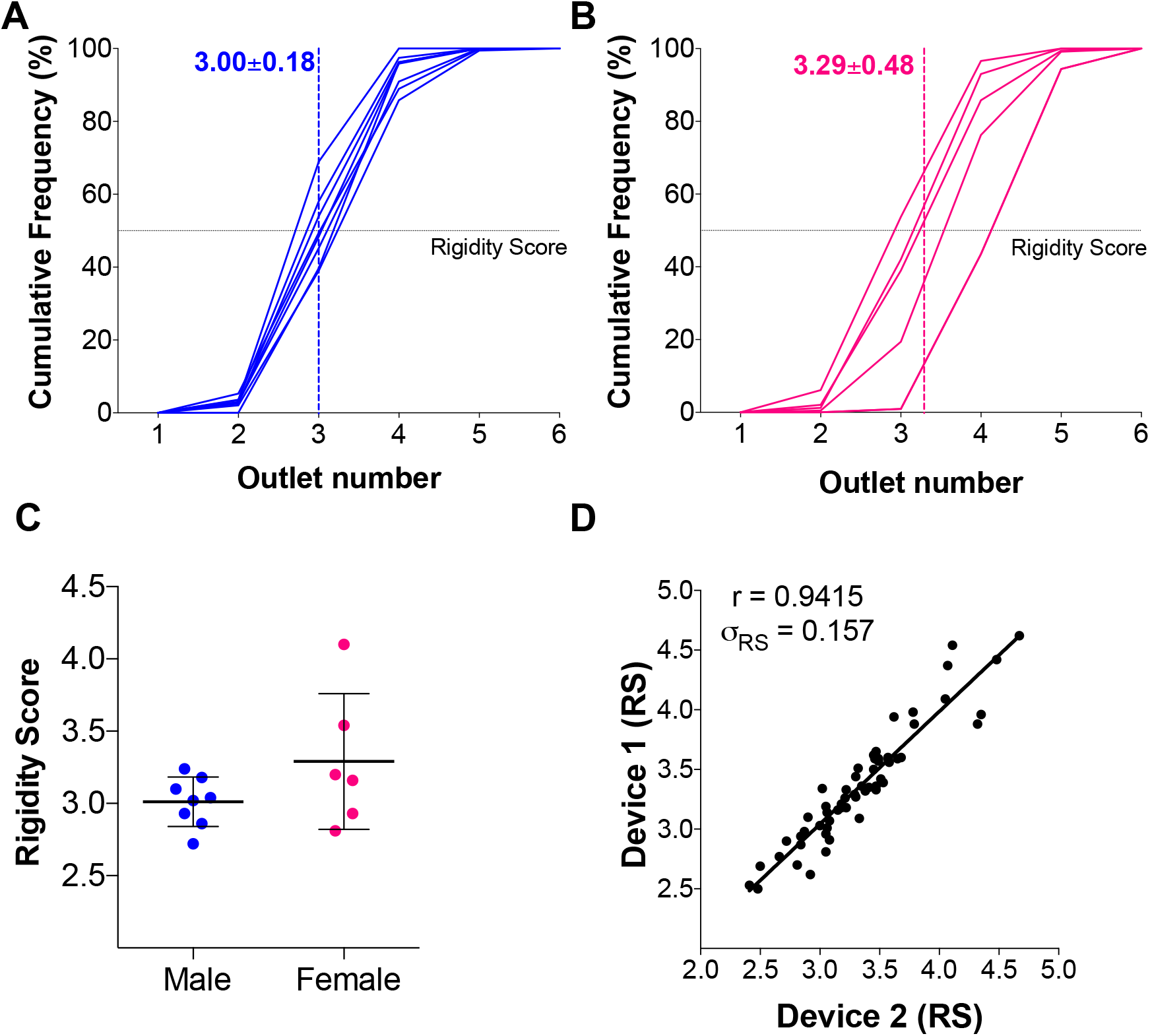
Inter-donor variability of packed RBC units on the day of collection and measurement repeatability. Cumulative distribution of RBCs sorted based on deformability using the microfluidic ratchet device from male (A) and female (B) donors. A rigidity score (RS) is derived from the fractional outlet number at the 50% cross over point of the cumulative distribution. The mean RS for male and female donors are indicated using dashed lines. (C) RS for RBCs from the day of collection for male (n=8) and female (n=6) donors. (D) Repeatability of the RS from doublet measurements on the same samples, which showed a Pearson’s r = 0.9415 and a standard deviation σ_RS_=0.157.

### RBC deformability profiles at time of collection

We established a baseline deformability profile of all blood bags at the time of collection. Packed RBCs in standard blood bags collected from healthy donors (n=14), were obtained from Canadian Blood Services. Consistent with previous reports^5,36^, we observed significant variability in initial RBC deformability among donors. Specifically, donor RS ranged from 2.71 to 4.13 with a mean of 3.13±0.37. The mean RS of male donor RBCs (n=8, 3.00±0.18, **Fig. 1A**) was slightly lower than female donor RBCs (n=6, 3.29±0.48, **Fig. 1B**), but this difference was not statistically significant (**Fig. 1C**).

### Measurement repeatability

Each RBC sample in this study was measured twice using different replicate microfluidic devices. We used this doublet data to confirm the repeatability of our measurement by plotting the RS from the first and second measurements against each other (**Fig. 1D**). These results suggest that the RBC deformability measurements were highly repeatable with a standard deviation of 0.157 in repeated measurements.

### RBC deformability loss during cold storage

To assess RBC deformability loss during cold storage, we sampled RBCs from blood bags every two weeks from 0 to 8 weeks of cold storage, which is two weeks beyond the current 42-day storage window (**Fig. 2**). From 0-4 weeks, the stored RBC units showed no detectable deformability loss. At the expiry date of 6 weeks, the stored RBC units showed detectable loss in deformability, as reflected by an increased RS of 0.35 (p<0.05; **Fig. 3A**). From weeks 6-8, the stored RBC units exhibited a dramatic loss of deformability. In fact, the average RBC deformability loss was greater from weeks 6-8 (ΔRS=0.42) than for the loss from weeks 0-6. While it should be noted that RBC deformability alone does not necessarily predict transfusion efficacy, this punctuated loss of RBC deformability after 6 weeks strongly supports the current 42-day storage window. Interestingly, some donated RBC units showed an initial increase in deformability from weeks 0-2, which was followed by a progressive deformability loss thereafter. These results are confirmed by doublet measurements and are also consistent with earlier studies that showed RBC units can often recover some of their deformability upon initial storage^5,47,48^.

**Fig. 2.**
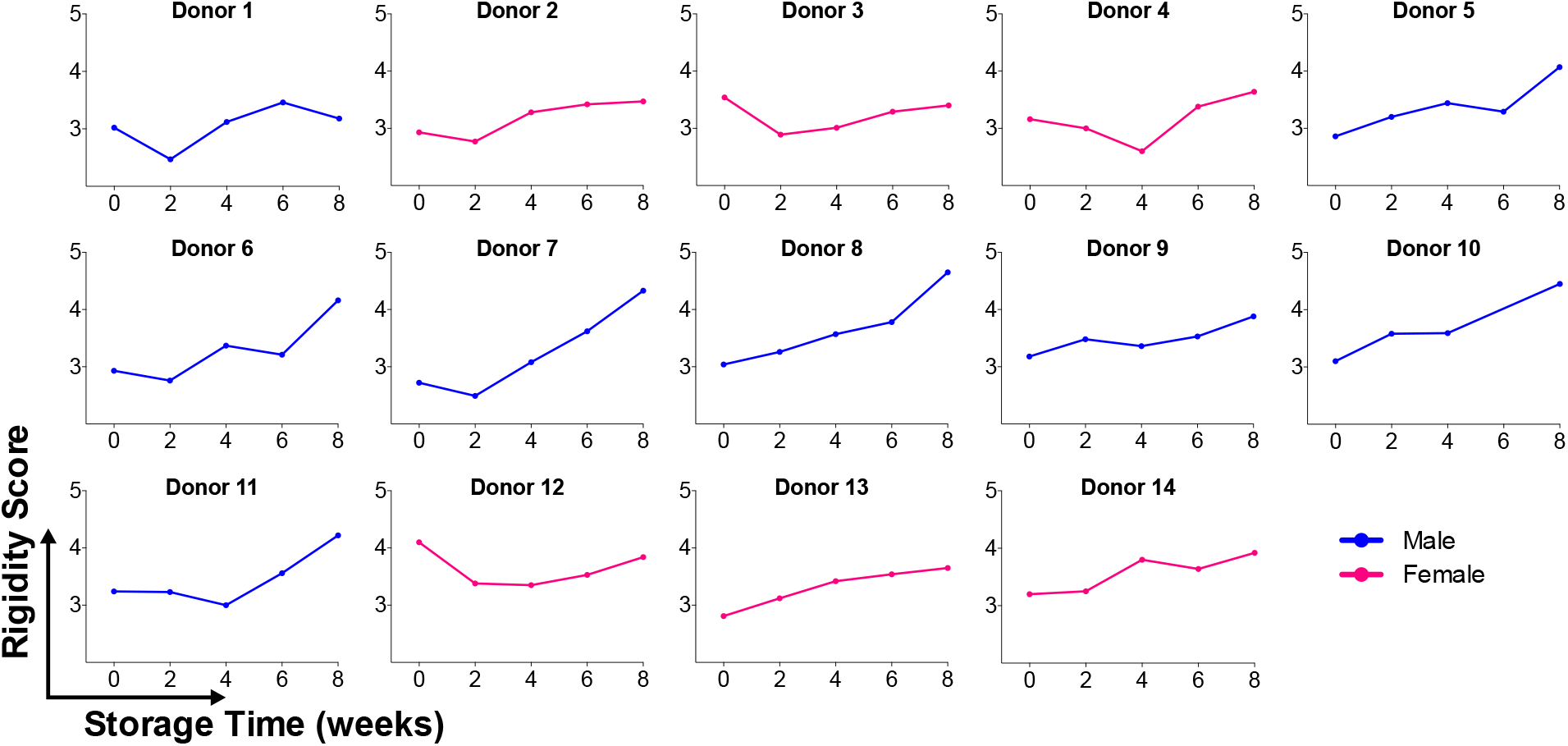
RBC deformability aging curves. Measured rigidity score (RS) for RBCs from each donor, sampled every two weeks over 8 weeks of cold storage. Each data point is the mean of doublet measurements.

We further evaluated whether the deformability of fresh RBC is predictive of the rate of RBC deformability loss during storage by relating the RS of RBCs at the time of collection to the RS at the end of the 42-day storage window. We found no correlation between the two (**Fig. 3B**). In fact, regardless of initial RS, all RBC units converged to RS of 3.48±0.16. These results confirm that the RBC deformability loss profile is cannot be predicted by initial RBC deformability and that determining this profile for each donor will require multiple samples over storage time.

**Fig. 3.**
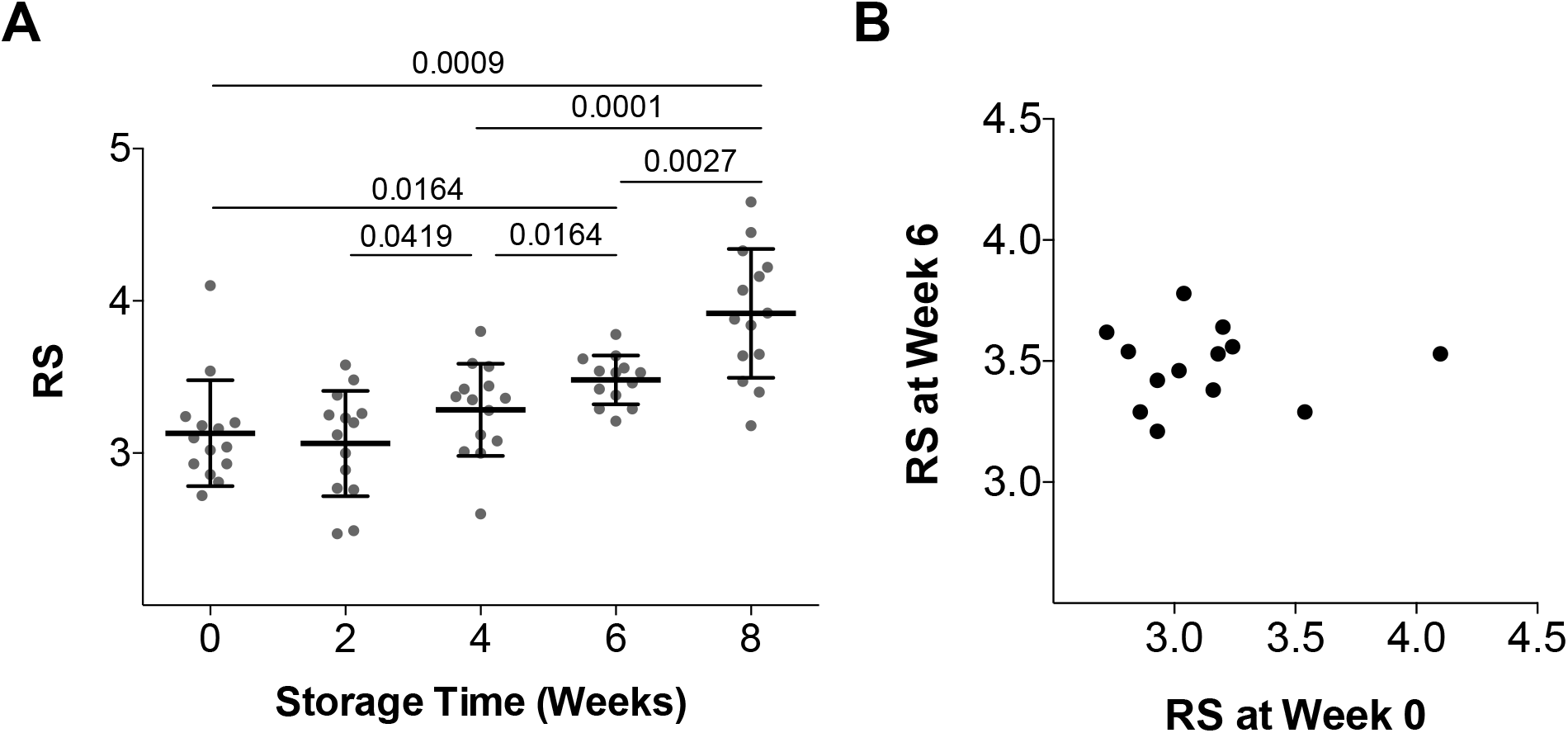
Degradation of RBC deformability during cold storage. (A) Donor RBCs exhibited a progressive increase in RS over 6 weeks, with an accelerated increase in RS between weeks 6-8. The mean rigidity scores (RS) were 3.13±0.35 (week 0), 3.06±0.35 (week 2), 3.29±0.30 (week 4), 3.48±0.16 (week 6), and 3.91±0.42 (week 8). (B) Correlation between RS on the day of processing and day of expiration (week 6), r=0.0086.

### Differences in RBC stability during storage between Male and Female donors

We investigated the differences in RBC deformability loss profiles between male donors (n = 8) and female donors (n = 6). In the first six weeks of storage, we observed no donor-specific variation in RBC deformability loss. From weeks 6-8, we observed a dramatic loss of RBC deformability for the male donors, but not female donors (p<0.05; **Fig. 4**). In fact, an accelerated RBC deformability loss in the final two weeks of storage was observed for the majority of the male donors (ΔRS=0.485) and for none of the female donors (ΔRS=0.172). These results suggest that certain donors are able to provide more stable RBCs and that these donors are more likely to be female than male.

**Fig. 4.**
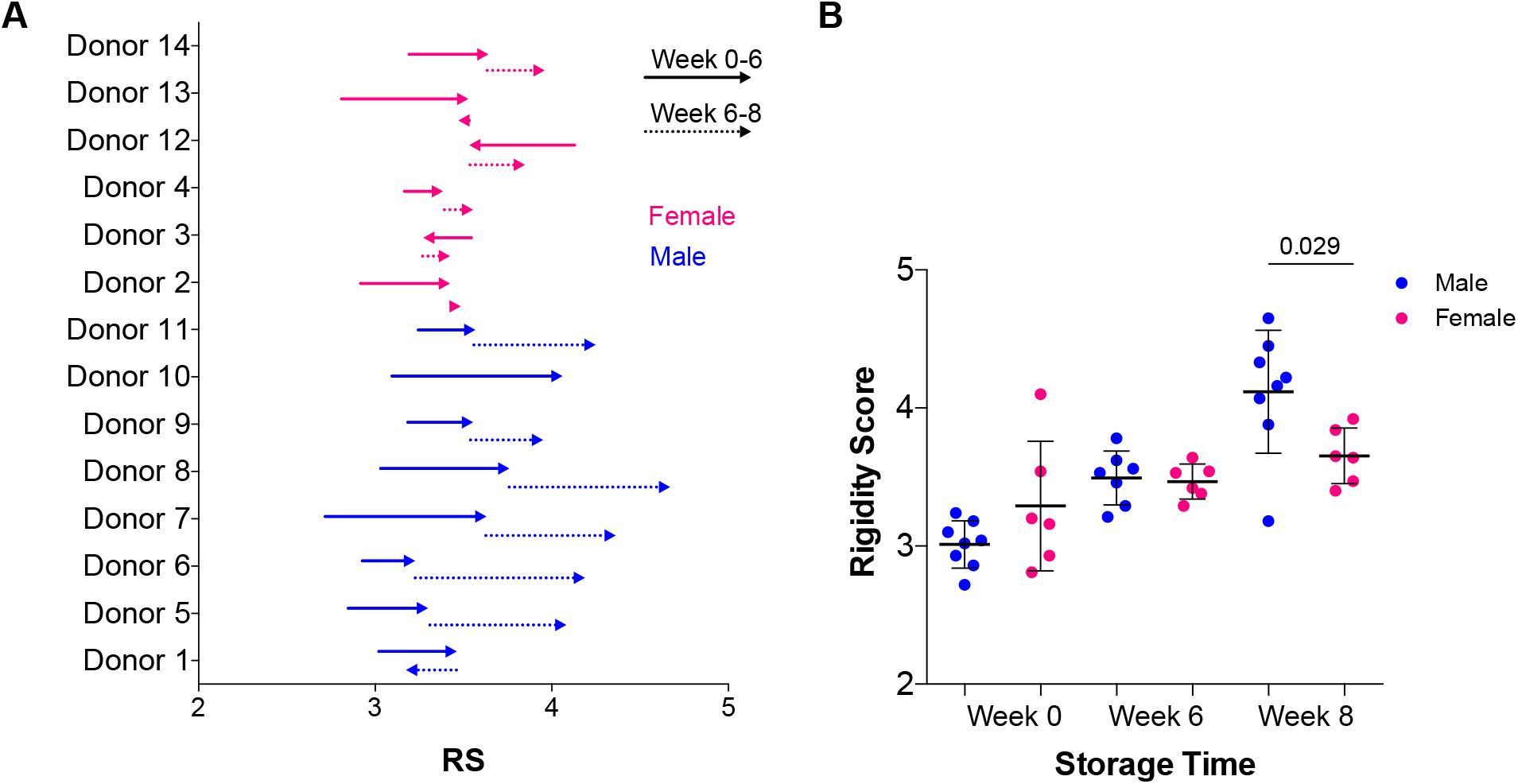
Comparison of RBC deformability loss profiles between male and female donors. (A) Changes in rigidity scores (RS) from weeks 0-6 and weeks 6-8 of cold storage. Arrows indicate direction of change. (B) RS for male and female donors at week 0, 6, and 8.

### Hematological parameters over 6 weeks of storage

We monitored standard hematological parameters (**Fig. 5**) of cold stored RBC units including mean corpuscular volume (MCV), red cell distribution width (RDW), mean cell hemoglobin (MCH) and mean corpuscular hemoglobin concentration (MCHC). Overall, the hematological parameters stayed within the normal range (**Fig. 5**, grey shaded area), with the notable exception of MCHC levels, which dropped slightly below accepted values of 315-355 g/L (Medical Council of Canada reference values^49^) at the 6-week expiration date. We related the general hematological data for each sample to the deformability of the matching RBCs at Week 0 and at Week 6 of storage. We found a slight positive correlation between increase in MCV and increase in the RS over time for male donors (r = 0.7504), but not for female donors. Male donors also showed a slight negative correlation between RS and MCHC (r = −0.6373). There were no correlations between any other parameters and changes in deformability.

**Fig. 5.**
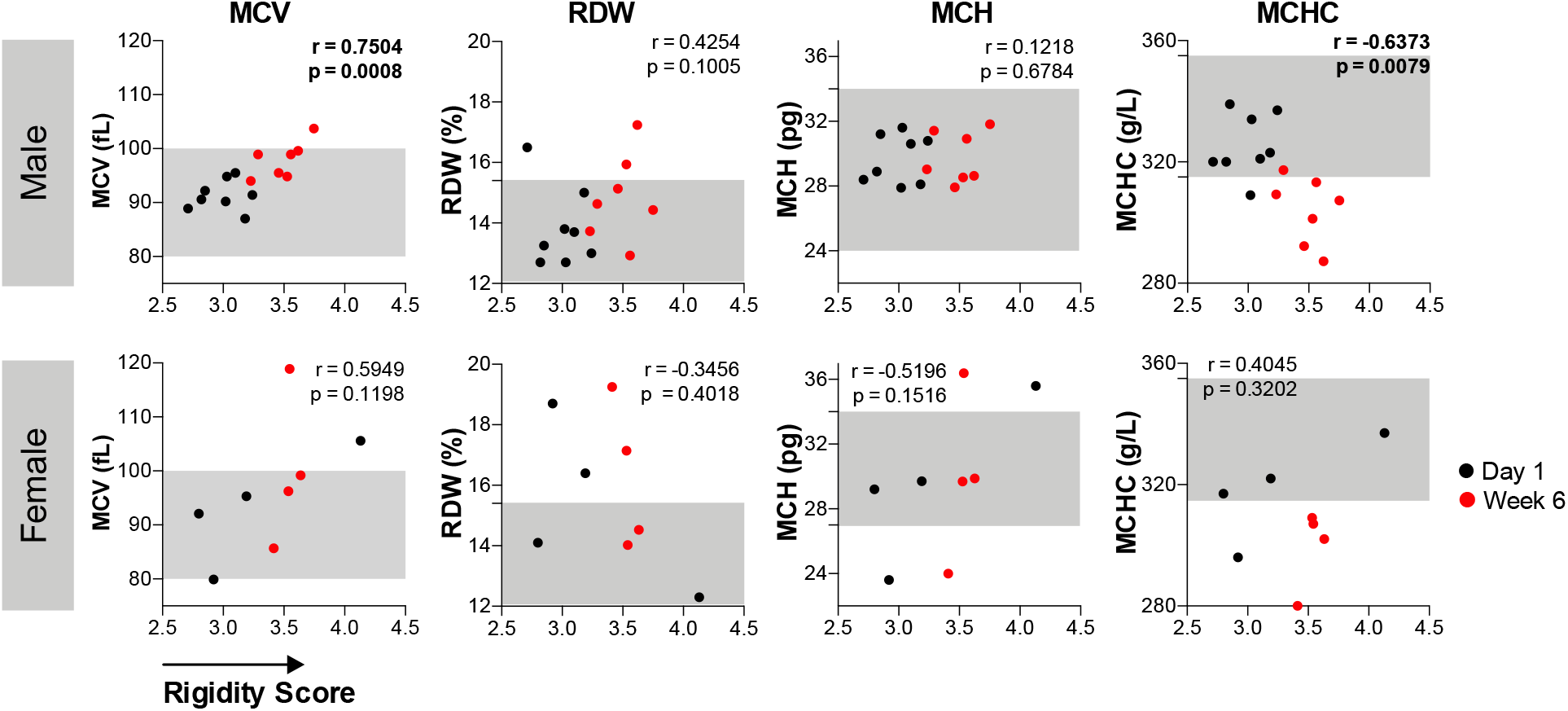
Correlation between hematological parameters and rigidity score (RS). Male donors showed minor correlation between RS and MCV (r=0.7504; p=0.0008), as well as between RS and MCHC (r=−0.6373; p=0.0079). Female donors showed no correlations between RS and hematological parameter.

### Hemolysis levels in blood bags

We also measured hemolysis at all time points for stored RBC units (**Fig. 6**). In Canada, the maximum allowable hemolysis is 0.8% at the time of expiry (6 weeks). We found that the majority of RBC samples did not show hemolysis above the standard safety level of 0.8% until week 8. The exceptions were Donor 9, which had 1.15% hemolysis at week 0, and Donor 13, which had 1.11% hemolysis at week 6. By week 8, half of the donor bags (n=3 male, n=2 female) were above the acceptable hemolysis threshold of 0.8%.

**Fig. 6.**
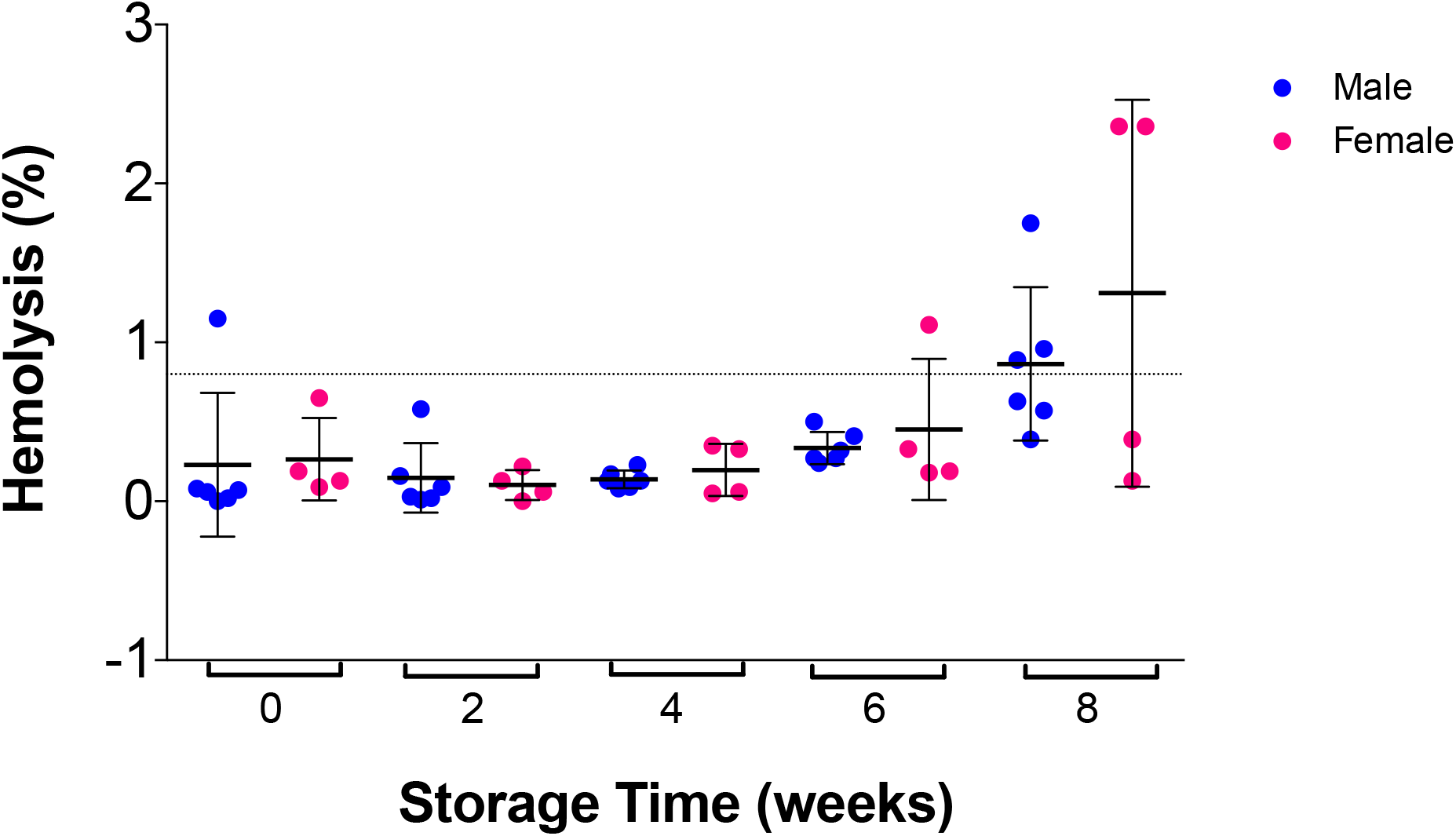
Hemolysis during storage in bags for male and female donors. The majority of donor blood bags did not show hemolysis above the 0.8% threshold, except for one male donor at week 0 and female donor at week 6. Half of all the blood bags (n=3 male and n=2 female) showed hemolysis above the 0.8% threshold at 8 week.

## DISCUSSION

In this study, we investigated the progressive loss of RBCs deformability under standard cold storage conditions. Using the microfluidic ratchet device, we sorted RBCs into fractions based on deformability and derived a RS based on the distribution of RBCs within these fractions. RS were obtained in freshly donated blood bags and over eight weeks of storage, which is two weeks longer than the standard storage window. We observed consistent loss of RBCs deformability during storage but the rate and magnitude of this loss was donor-specific and was not predicted based on the deformability of freshly donated RBCs. After 6 weeks of storage, RBCs from both male and female donors converged to a similar deformability. However, from weeks 6-8, the RBC deformability loss accelerated dramatically for male donors, but not for female donors. Together, these results demonstrate how RBC deformability could be used as a potential biomarker for the storage lesion on donated RBC units, and that a different storage window may be appropriate for certain donors.

The observed differences between male and female donors in their RBC deformability loss profiles is consistent with other differences between male and female blood. For example, RBCs from males have been shown to be smaller in size, as well as greater in hematocrit, MCV, hemoglobin concentration, viscosity and RBC fragility compared to RBCs from females^42^. Some of these differences have been attributed to the female sex hormone estrogen, which has been shown to protect RBCs from deformability loss^43^, but also has a major impact on the regulation of erythropoiesis^44^. Furthermore, difference in distribution and function of estrogen receptors on RBCs^45^, as well as differences in serum estradiol concentration may^46^ affect intracellular signalling and better protect against oxidative stress in female RBCs. These differences could collectively explain the accelerated deformability loss observed for RBCs from male donors upon storage past the 42-day storage window.

Profiling loss of RBC deformability for individual donors could serve to guide the selection of blood units prior to transfusion. It is well-established that clinical efficacy of RBC units in blood transfusions declines with the age of the blood bag^33–35,37,38^ and that the loss of RBC deformability corresponds with this decline in clinical efficacy^47^. This study demonstrates that loss of RBC deformability can be profiled over the course of storage. We observed that RBC deformability was generally preserved during the 42-day storage expiration window. However, beyond the 42-day storage expiration window, RBC deformability loss was accelerated and varied significantly between donors. Donor-specific variability in deformability loss of stored RBCs is consistent with previous reports^5,36,39^ and may strongly impact post-transfusion outcomes^32,50,51^. Consequently, RBC deformability profiling could be a valuable tool to screen for donors whose RBCs could be reliably stored for longer than the 42-day storage window.

## Data availability statement

The data that support the findings of this study are available on request from the corresponding author.

## Ethics approval statement

This study was approved by the University of British Columbia’s Clinical Research Ethics Board (UBC REB# H19-01121) and Canadian Blood Services Research Ethics Board (CBS REB# 2019-029).

## Patient consent statement

Patient consent was not mandated for this study.

## Clinical Trial Registration

This study was not a clinical trial.

## Acknowledgments

We are grateful to Canadian Blood Services’ blood donors who made this research possible. We would also like to thank Dana Devine and Brankica Culibrk for providing access and training for the Sysmex system.

## Author contributions

H.M. supervised the study. H.M., E.I., and K.M. conceived the idea. E.I., K.M., and E.L. performed the experimental work. All authors wrote the manuscript.

## SUPPLEMENTALS

**Figure S1.**
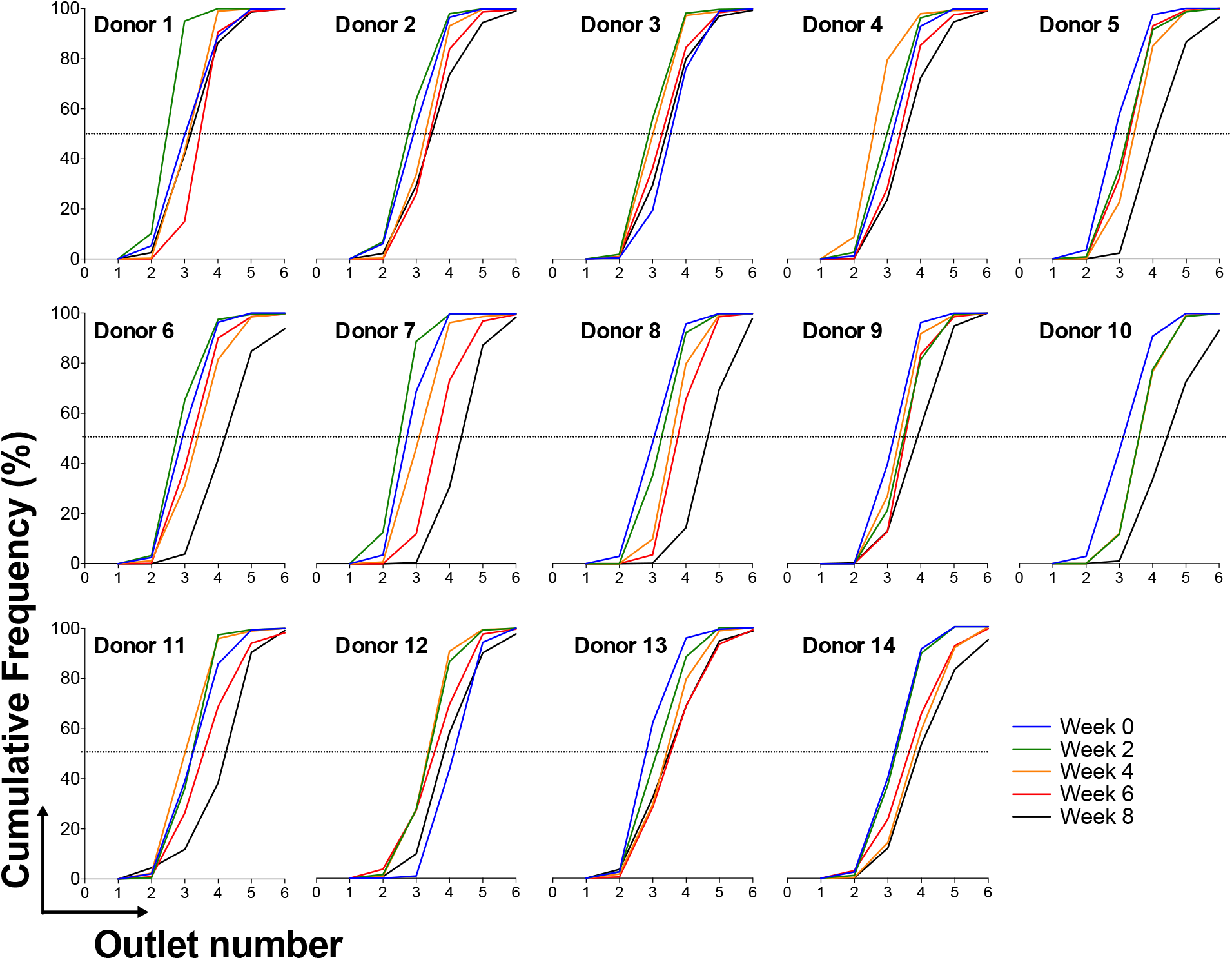
Cumulative distribution curves after deformability-based sorting of RBCs using the microfluidic ratchet device. (A) Cumulative distribution curves from deformability sorting of RBC units at Week 0 (day of manufacturing, blue line), followed by Weeks 2 (green), 4 (orange), 6 (red), and 8 (black) of cold storage. Each donor showed distinct RBC deformability loss profiles (shift to the right) during storage.

